# Photodynamic priming with Vitamin D and ALA-based PDT induces intratumoral immune cell recruitment and signaling pathway activation in murine cutaneous squamous cell carcinoma

**DOI:** 10.64898/2026.02.04.703788

**Authors:** Alan S. Shen, Sanjay Anand, Cheng-En Cheng, Benjamin Kovacic, Jennifer Powers, C. Marcela Diaz-Montero, Tayyaba Hasan, Edward V. Maytin

**Author notes:** These authors contributed equally to this work (Alan S. Shen, Sanjay Anand). Corresponding Author Information: Edward Maytin, MD PhD Mailstop ND-20, Cleveland Clinic Research Cleveland Clinic 9500 Euclid Avenue Cleveland, OH 44195.

## Abstract

Photodynamic therapy (PDT) is effective for early epithelial pre-cancers, yet its efficacy in fully-developed cutaneous squamous cell carcinoma (SCC) is limited by immunosuppression in the tumor microenvironment. Vitamin D (VitD) has emerged as a potential neoadjuvant to enhance photodynamic priming in several cancers. Here, we investigated how VitD pretreatment modulates local and systemic immune responses to aminolevulinic acid–based PDT in two immunocompetent murine SCC models (chronic UV-induced SCC, and subcutaneously implanted PDVC57B cells). VitD combined with PDT significantly amplified hallmarks of immunogenic cell death, including calreticulin and HMGB1 expression, and increased recruitment of neutrophils, macrophages, dendritic cells, and CD8⁺ T cells into tumors. Importantly, VitD+PDT produced a higher M1/M2 macrophage ratio, and reduced the number of exhausted (PD-1 expressing) T cells compared to PDT alone. Immune profiling of blood demonstrated enhanced T-cell activation (CD69) and reduced TIM-3 expression on cytotoxic T cells. Transcriptomic analysis revealed pathway enrichment for interferon-α/γ signaling and suppression of pro-tumorigenic epithelial–mesenchymal transition and of angiogenesis after VitD+PDT. Collectively, these findings demonstrate that VitD reprograms the immune response to PDT by enhancing cytotoxic immunity while limiting immunosuppressive features. This suggests an immune-priming strategy to consider, possibly alongside immune checkpoint blockade, for treating cutaneous SCC.

## INTRODUCTION

Non-melanoma skin cancers (NMSC), including squamous cell carcinoma (SCC) and basal cell carcinoma (BCC), are the most common of all human malignancies[1][2][3]. Surgical excision is the first-line treatment for SCC to prevent local invasion and metastases[4, 5][6]. Although locally advanced disease and metastases are relatively uncommon (up to 10% and 1% of cases, respectively), the absolute number of cases is significant and systemic therapy is often required[6]. Immune checkpoint inhibitors (ICIs) have shown promise in advanced NMSC, but only a limited subset of patients respond[4]. Therefore, additional effective therapies and adjuvants for SCC are needed.

Photodynamic therapy (PDT) is a non-invasive, non-scarring, and non-mutagenic treatment for skin cancers and precancers. A precursor drug (5-aminolevulinic acid; ALA) is preferentially taken up by tumor cells and converted into a photosensitizer, protoporphyrin IX (PpIX). Upon activation with visible light, PpIX generates reactive oxygen species that induce tumor cell death[7]. Direct killing of tumor cells by PDT is well-known, but the concept of *photodynamic priming* (*PDP*) is relatively new[8, 9]. PDP refers to a collection of PDT-induced mechanisms that occur in surviving but damaged tumor tissue, which together act to enhance the therapeutic response to additional therapies (e.g., chemotherapy or ICIs)[8, 9]. PDP effects include loosening of tumor stroma, induction of immunogenic cytokines, and triggering of immunogenic cell death (ICD). ICD is characterized by expression of damage-associated molecular patterns (DAMPs) that promote recruitment and activation of neutrophils, macrophages, and dendritic cells[10, 11]; these innate immune cells then drive a subsequent, adaptive immune response against the tumor.

To futher enhance PDP effects, *Vitamin D* (VitD) has emerged as a promising neoadjuvant due to several complementary actions. First, VitD promotes squamous cell differentiation and increases intracellular PpIX accumulation[12]. Second, VitD can regulate innate and adaptive immunity in the setting of cancer, through VitD receptor–mediated gene expression[13][14]. Third, a randomized clinical trial in patients with actinic keratoses (AK; squamous pre-cancer) confirmed that high-dose VitD supplementation prior to PDT significantly improves lesion clearance rates[15]. A different clinical trial found that high-dose VitD and PDT can enhance lesion clearance in patients with BCC, particularly for thin BCC tumors[16]. While such preclinical and clinical results highlight the potential utility of VitD+PDT for patients with AK and thin BCC, the question of whether these immune-enhancing mechanisms are operative in fully-developed SCC remained unanswered. In this study, we used two immunocompetent murine SCC models to characterize the magnitude, time-course, and functional significance of immune changes induced by VitD+PDT compared with each treatment alone.

## METHODS

### Cell Lines and Culture

Murine PDVC57B (PDV) squamous cell carcinoma cells (CancerTools[17]) were originally derived from C57BL/6 keratinocytes via repeated dimethylbenz(a)anthracene (DMBA) treatment[18]. Cells were cultured in Dulbecco’s Modified Eagle Medium (DMEM) supplemented with 10% fetal bovine serum and 1% penicillin–streptomycin at 37°C in 5% CO₂. Cultures were passaged every 2 days at a 1:3 ratio, and cells at ≥ 5 passages were used for in vivo experiments.

### UV-Induced SCC Model

Female SKH-1 hairless mice (8 weeks old) were exposed to UV irradiation (80% UVB/20% UVA) three times weekly, beginning at 80 mJ/cm² and increasing by 10 mJ/cm² weekly to a maximum of 180 mJ/cm². By 20 weeks, mice developed morphologic and histologic features consistent with human SCC. All animal protocols were approved by the Cleveland Clinic IACUC.

### Subcutaneous Syngeneic SCC Model

PDVC57B cells (2–5 × 10⁶) were resuspended in 0.1 mL of a 1:1 Matrigel/DMEM mixture and injected subcutaneously into the left flank of 8-week-old C57BL/6 mice. Tumors reached 5–10 mm in 2–3 weeks.

### Vitamin D Pretreatment

For superficial UV-SCC lesions, topical calcitriol ointment (3 μg/g; Vectical) or vehicle (petroleum jelly) was applied daily for 3 days (72 hr) prior to PDT[19]. For implanted PDV tumors, calcitriol (1 μg/kg) was administered intraperitoneally once daily for 3 days. An overview of treatment conditions and the samples collected is shown in **Figure 1**.

**FIG. 1.**
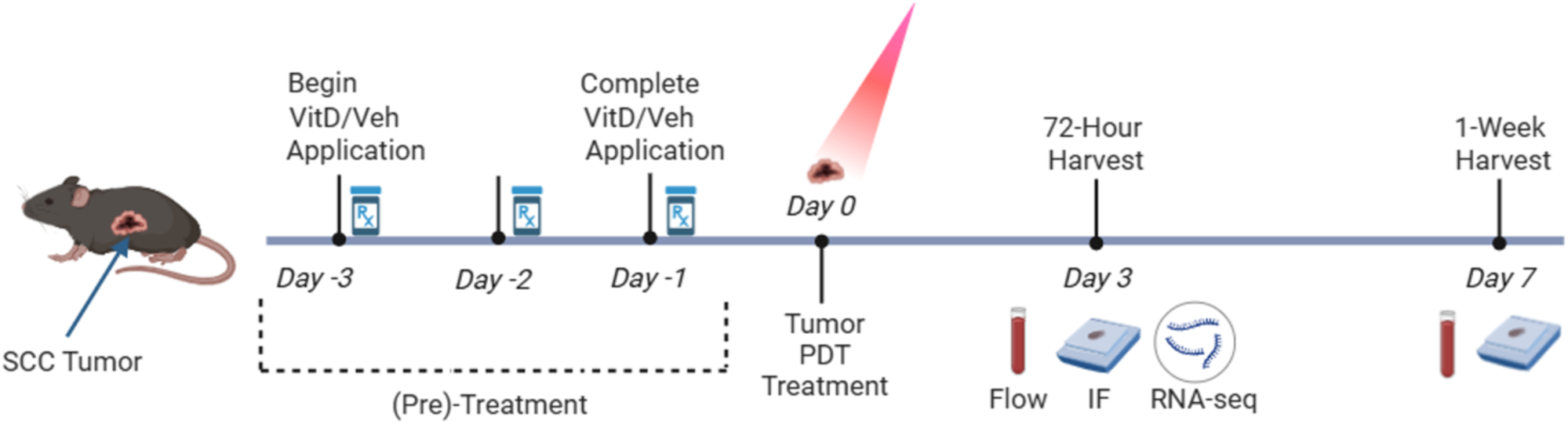
Experimental design for VitD pretreatment and PDT in murine SCC models. SCC tumors were generated using either a UV-induced model in SKH-1 hairless mice or a subcutaneous PDVC57B cell–derived model in C57BL/6 mice. VitD pretreatment was administered over 72 hours (Days –3 to –1), just prior to PDT on Day 0. Tumors receiving no PDT (VitD alone or vehicle control) were harvested after the same 72-hour pretreatment period. PDT-treated tumors were harvested at 72 hours (Day 3) or 1 week (Day 7) post-treatment. Peripheral blood was collected at each tissue harvest for flow cytometry, and tumor tissue was processed for immunofluorescence (both time points) and bulk RNA sequencing (72-hour time point only).

### Photodynamic Therapy

UV-SCCs received topical 20% 5-aminolevulinic acid (ALA) in PBS/EDTA/DMSO, followed immediately by blue light exposure (Blu-U; 417 nm, 30 min, 18 J/cm²). PDV tumors received intraperitoneal ALA (200 mg/kg) with a 4-h uptake period, followed by red light illumination (LumaCare xenon source; 633 nm, 100 J/cm²). Tumors were shielded with opaque foil templates to confine illumination[11, 19]. Blue light was used for superficial UV-SCCs; red light was selected for PDV tumors due to its deeper penetration of tissues.

### Tissue Collection and Immunofluorescence

Tumors were excised at 72 h or 1-week post-PDT, fixed in 10% neutral-buffered formalin overnight, and paraffin-embedded. Sections (5 µm) were deparaffinized, rehydrated, and subjected to citrate buffer antigen retrieval. After blocking with 3% normal donkey serum, slides were incubated overnight at 4°C with primary rabbit antibodies against Ly6G (neutrophils), F4/80 (macrophages), CD86 (M1 macrophages), CD206 (M2 macrophages), CD11c (dendritic cells), CD3 (T-cells), CD8 (cytotoxic T-cells), FoxP3 (regulatory T-cells), or PD-1 (exhaustion T-cells). All primary antibodies were from Cell Signaling Technology, Danvers, MA. Cy3-conjugated secondary antibodies (Jackson ImmunoResearch Laboratories, West Grove, PA) and DAPI mounting medium (VECTASHIELD) were used for detection. Three high-power fields per tumor were imaged by fluorescence microscopy. Immune cells were counted manually; DAMP expression was quantified by ImageJ[20].

### Peripheral Blood Flow Cytometry

Prior to tumor harvesting, peripheral blood was collected from each mouse via terminal cardiac puncture. Samples were first incubated with Fc block (anti-mouse CD16/CD32, Clone 93) to reduce non-specific antibody binding. Samples were then stained with fluorochrome-conjugated antibodies and analyzed on a BD FACSymphony A5 SORP flow cytometer.

Unless otherwise noted, antibodies for immune profiling were purchased from BD Biosciences and targeted the following: CD45 (Clone 30-F11), Ly6G (1A8), CD11c (HL3), NK1.1 (PK136), CD8 (53–6.7), CD11b (M1/70), CD3e (500A2), CD335/NKp46 (29A1.4), and Ly6C (AL-21). For T regulatory cell identification: CD4 (GK1.5), CD25 (PC61), and CD127 (SB/199). For immune activation and functional classification: CD154 (MR1), CD44 (IM7), CD62L (MEL-14), CD69 (H1.2F3), I-A/I-E (BioLegend; M5/114.15.2), CCR2 (475301), and CXCR2 (V48–2310). For immune checkpoint markers: PD-1 (J43), PD-L1 (eBioscience; MIH5), ICOS/CD278 (BioLegend; 15F9), and TIM-3 (BioLegend; B8.2C12).

Flow cytometry data were analyzed using FlowJo software (version 10.8; Treestar, Ashland, OR, USA). The gating scheme utilized for each sample has been previously published[19].

### RNA Isolation and Sequencing

A portion of each tumor (≤30 mg) was stored in RNAlater, frozen, and processed (Qiagen RNeasy Mini Kit). RNA concentration and integrity (DV200) were confirmed, and libraries were prepared using Illumina Stranded mRNA Prep with poly(A) selection. Sequencing was performed on an Illumina NovaSeq 6000 (≈30 million paired-end reads/sample).

Raw sequencing reads in FASTQ format were aligned to the latest mouse reference genome (GRCm39) using the STAR aligner (version 2.7.11a)[21]. Alignment quality control was conducted using RSeQC, which assesses mapping quality and RNA integrity metrics from the resulting BAM files[22]. Read summarization was carried out using featureCounts[23], which quantified the number of reads mapped to each gene based on the GENECODE M33 mouse annotation (GRCm39). Gene-level count data from all samples were merged using an in-house R script. Differential gene expression analysis was performed using the DESeq2 package[24]. DESeq2 normalizes read counts to account for differences in sequencing depth and library composition using the median-of-ratios method. It models gene expression variation using a negative binomial distribution and estimates gene-specific dispersion to account for heteroscedasticity. Statistical significance for differential expression between groups was assessed using the Wald test, and p-values were adjusted for multiple hypothesis testing using the Benjamini-Hochberg procedure to control the false discovery rate. The final DESeq2 output included a list of differentially expressed genes (DEGs) with associated log2 fold changes and adjusted p-values (FDRs).

Gene Set Enrichment Analysis (GSEA) was then performed on the ranked list of DEGs using the clusterProfiler R package[25]. Enrichment was tested against the Hallmark gene sets from the Molecular Signatures Database (MSigDB), which represents 50 curated biological pathways[26]. Visualization of enriched pathways was performed using the dotplot function from the enrichplot R package[27], which displays the enrichment score and adjusted p-values across all pathways.

### Statistics

A power analysis was performed based on the primary endpoint of neutrophil proportion following either photodynamic therapy (PDT) alone or vitamin D pretreatment plus PDT. Mean and standard deviation estimates were derived from preliminary data on neutrophil responses to PDT in murine actinic keratoses[19]. To detect a 15% difference in group means with 80% power and a significance level of 0.05, a minimum of 4 mice per treatment group was determined to be sufficient.

Data from immunofluorescence cell counts and flow cytometry–derived cellular proportions were analyzed in GraphPad Prism v10.0 (GraphPad Software, San Diego, CA). Group differences were assessed using the nonparametric Kruskal–Wallis test, followed, when significant, by pairwise Mann–Whitney U tests. These pairwise Mann–Whitney comparisons were prespecified (Veh vs. VitD, Veh vs. VehPDT 72hr, Veh vs. VitD+PDT 72hr, Veh vs. Veh+PDT 1wk, Veh vs. VitD+PDT 1wk, Veh+PDT 72hr vs. VitD+PDT 72hr, Veh+PDT 1week vs. VitD+PDT 1week). To control for multiple comparisons testing, raw p-values from these pairwise comparisons were adjusted using the Benjamini–Hochberg false discovery rate (FDR) procedure. Adjusted p values were reported using GraphPad’s default significance notation, where p < 0.05 was denoted by one asterisk (*), p < 0.01 by two asterisks (**), p < 0.001 by three asterisks (***), and p < 0.0001 by four asterisks (****); p ≥ 0.05 was considered not significant (“ns”). Data are presented as mean ± SEM unless otherwise noted. Significance bars are shown for Veh vs other groups (only the first significant comparison displayed for IF data), as well as for Veh + PDT 72hr vs VitD+PDT 72hr and Veh+PDT 1wk vs VitD+PDT 1wk.

For comparisons with n = 3 per group (M1 macrophages, M2 macrophages, PDV IF data), statistical power was insufficient to detect significance reliably; therefore, these results are presented without significance bars and are interpreted as descriptive trends in the data.

## RESULTS

To evaluate how the anti-tumor immune response to PDT is altered when combined with neoadjuvant VitD, we used two different tumor models, one created by chronic UV exposure (UV-SCC) and the other by subcutaneous injection of an SCC cell line (PDVC57B). Each model was used in time-course experiments outlined in **Figure 1**.

### Vitamin D and PDT-induced changes in expression of damage associated molecular patterns (DAMPs) in UV-SCC tumors

To assess the impact of VitD and PDT on DAMPs, we performed immunofluorescence analysis (IF) of calreticulin and high mobility group box 1 (HMGB1) expression in UV-SCC tumors. Compared with vehicle, VitD, PDT (72 hr post-treatment), and VitD+PDT (72 hr) each significantly increased calreticulin (p < 0.01, **Figure 2A**) and HMGB1 (p < 0.01, **Figure 2B**). Notably, VitD+PDT at 72 hr produced significantly higher levels of both calreticulin and HMGB1 than PDT alone (p < 0.01) at the same time point.

**FIG. 2.**
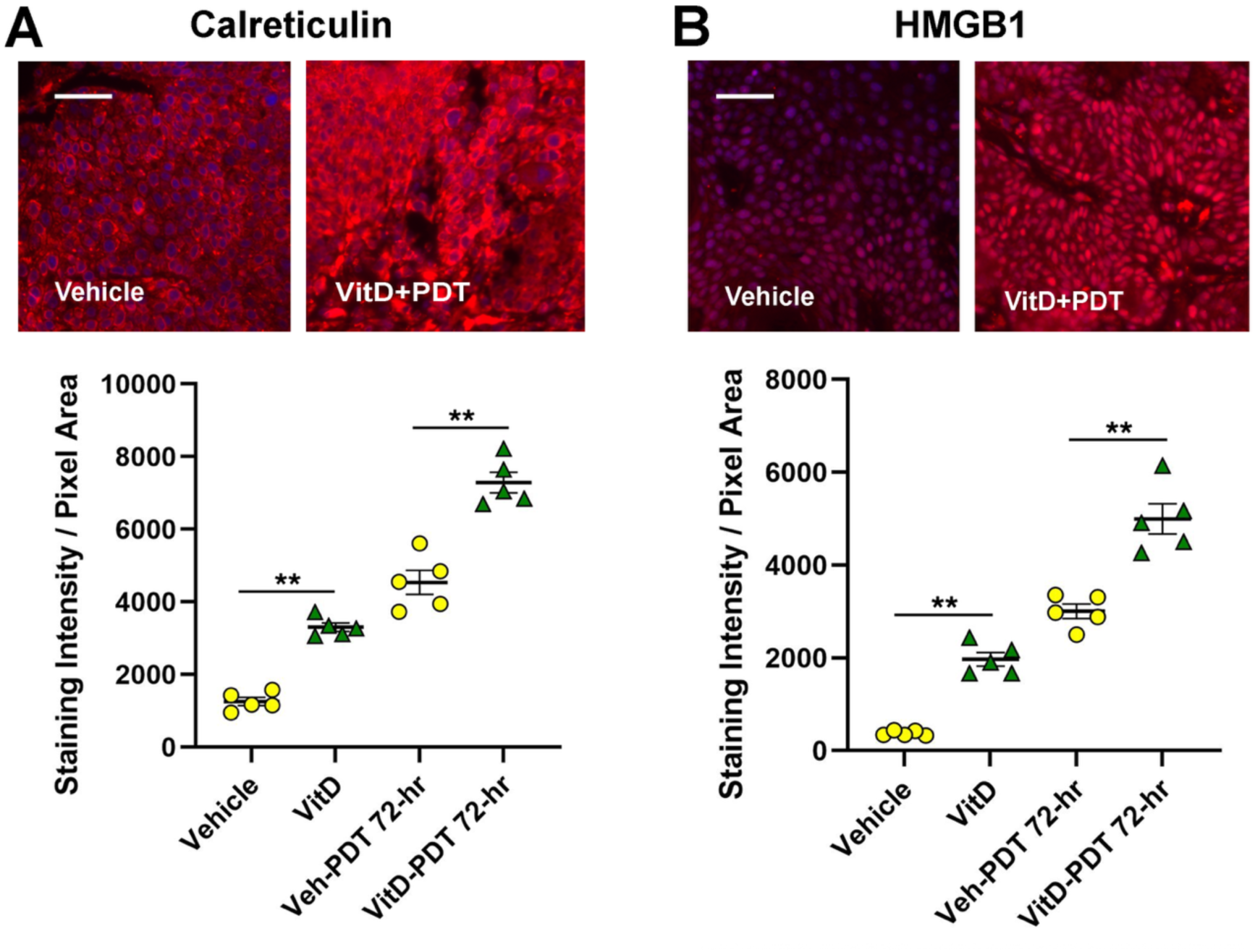
Induced expression of the damage-associated molecular pattern (DAMP) proteins *calreticulin* and *HMGB1* in UV-induced SCC lesions following VitD ± PDT. For (**A**) calreticulin and (**B**) HMGB1, representative immunofluorescent images are displayed above summary graphs that quantify the staining intensity per unit area (pixel) in lesional tissue harvested either before PDT or 72 hours after VitD ± PDT. *n* = 5 mice per group; each point represents the average of three fields per lesion from each mouse, with mean ± SEM displayed. Statistical analysis used Kruskal–Wallis for overall group comparisons, and when significant, Mann–Whitney U-tests for pairwise comparisons with adjustment for multiple comparisons utilizing the Benjamini-Hochberg correction. Significance levels: *(**), p < 0.01.* Scale bars = 100 μm.

### Vitamin D and PDT-induced changes in recruitment of innate immune cells in UV-SCC tumors

Next, to determine how VitD and PDT influence innate immune cell recruitment in UV-SCC tumors, we quantified neutrophils (Ly6G+), dendritic cells (CD11c+), and macrophages (F4/80+). The number of neutrophils was significantly increased after VitD alone, PDT alone (72 hr), and VitD+PDT (72 hr), when compared with vehicle (**Figure 3A**). VitD+PDT (72 hr) also led to significantly higher neutrophil levels than PDT alone (72 hr), indicating an additive effect of VitD (**Figure 3A**). Similar trends were observed for macrophages (**Figure 3B**), and for dendritic cells (**Figure 3C**), with effects persisting up to 1-week post-PDT, at least for macrophages.

**FIG. 3.**
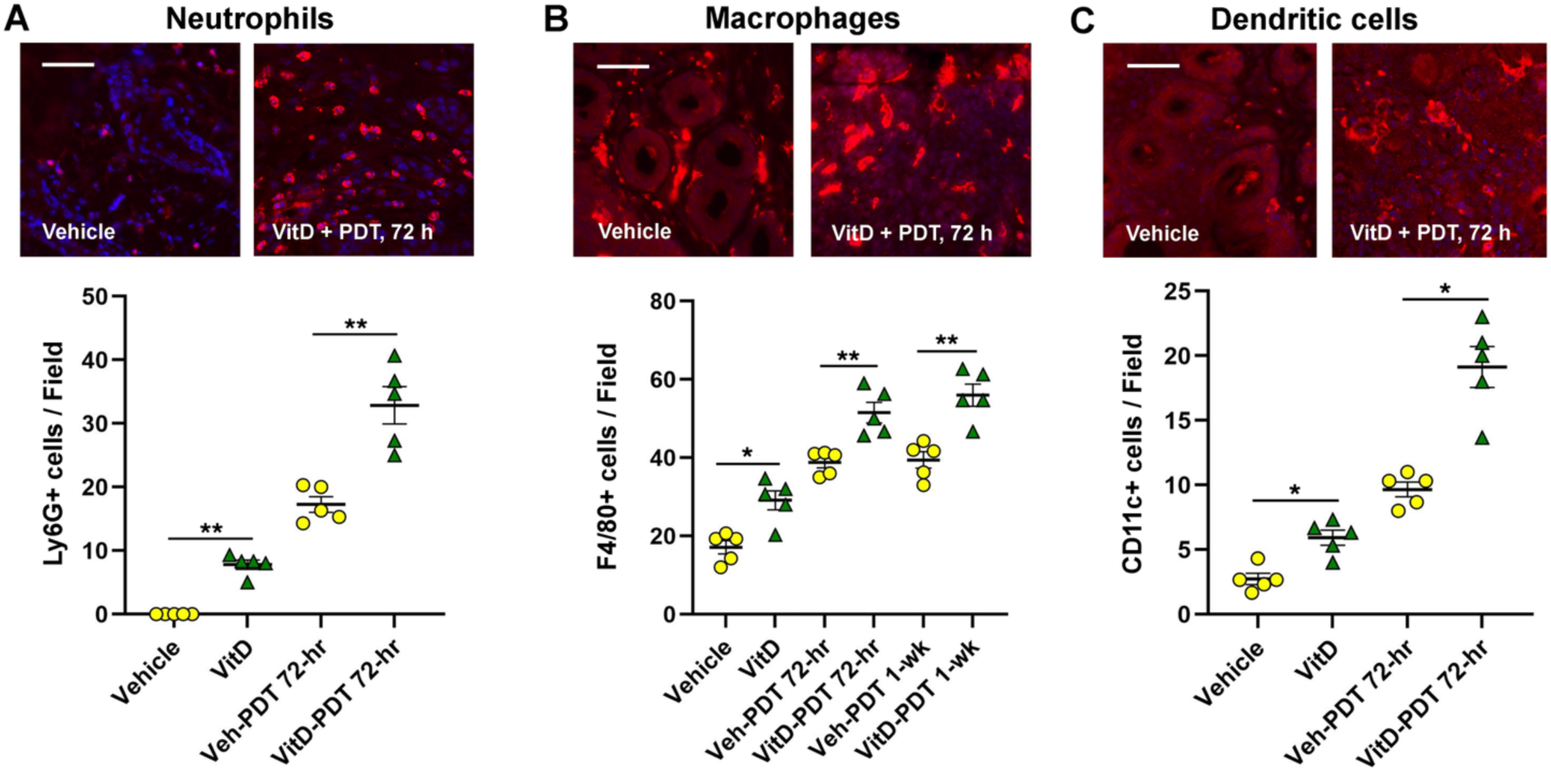
Recruitment of innate immune cells: Recruitment of neutrophils, macrophages, and dendritic cells into UV-induced SCC lesions following VitD ± PDT. Representative images of each type of immunostain, and complete quantification of cell numbers per field for: (**A**) neutrophils; (**B**) macrophages; and (**C**) dendritic cells in the SCC tumors at 72 hours after no treatment (Vehicle), VitD only, PDT only, or VitD+PDT are shown. Each point represents the average of three fields/lesion (one lesion/mouse); with *n* = 5 mice/group, there are 15 total data points/group, with mean ± SEM displayed. Statistical analysis was performed as in Figure 2. Significance levels: (\**), p* < 0.05; *(**), p < 0.01*. Scale bars = 100 μm.

### Vitamin D and PDT-induced changes in recruitment of adaptive immune cells in UV-SCC tumors

When assessing the effect of VitD and PDT on adaptive immune cell recruitment in UV-SCC tumors, total T cells (CD3+) were significantly increased after VitD alone, as well as after PDT alone or after VitD+PDT at 72 hr and at 1 week post-PDT, relative to vehicle-only controls (**Figure 4A**). An additive effect of the VitD+PDT combination was observed at both 72-hr and 1 week (p < 0.01). A similar pattern was observed for CD8+ T cells (**Figure 4B**). T-regs (FoxP3+) were unchanged with VitD alone but increased after PDT and VitD+PDT at 1 week, with higher levels after VitD+PDT than PDT alone (**Figure 4C**).

**FIG. 4.**
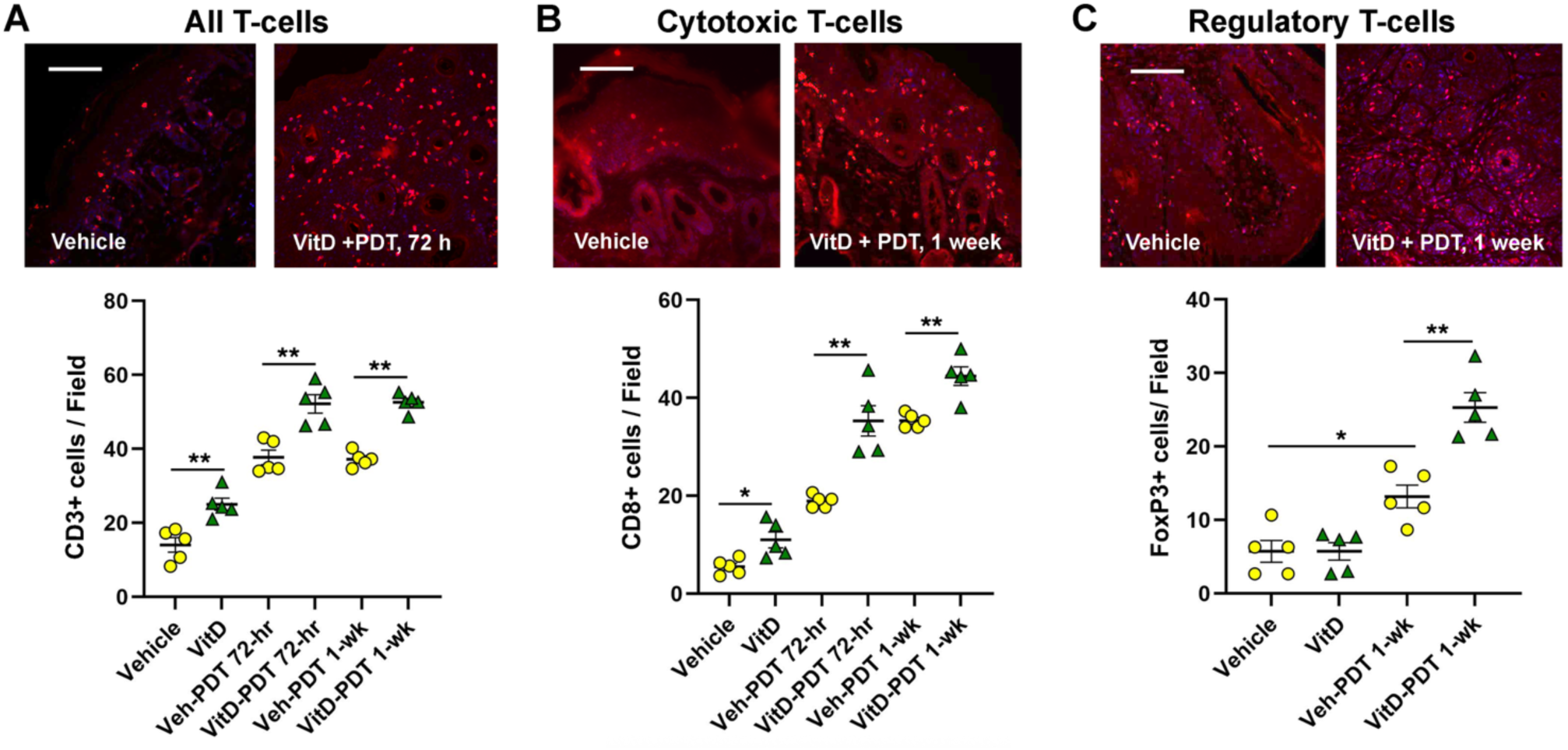
Recruitment of adaptive immune cells: T-cells, CD8+ T-cells, and T-regs in UV-induced SCC following VitD ± PDT. Quantification of: (**A**) total T-cells (CD3+), (**B**) CD8⁺ T-cells; and (**C**) T-regs per field is shown below representative immunofluorescently-stained images of these cells in UV-SCC lesions at 72 hours after VitD+PDT. Each point represents the average of three fields per lesion (one lesion/mouse); with *n* = 5 mice/group, for a total of 15 total data points/group; the mean ± SEM is displayed. Statistical analysis was performed as in Figure 2. Significance levels: (\**), p* < 0.05; *(**), p < 0.01*. Scale bars = 100 μm.

### Vitamin D and PDT-induced changes in immunocytes in the PDVC57B subcutaneous model

To further validate immune cell recruitment patterns in SCC tumors, we examined the second tumor model. In PDVC57B tumors, VitD or PDT alone increased neutrophils, macrophages, and dendritic cells, with greater increases from VitD+PDT (**Figure 5A–C**). VitD and PDT each elevated total T cells and CD8⁺ T cells, further augmented by VitD+PDT (**Figure 5D, E**). T-regs increased with PDT and VitD+PDT but not VitD alone (**Figure 5F**), which was an observation similar to the UV-SCC model. PD-1⁺ cells were decreased with VitD, increased with PDT, and returned toward baseline with VitD+PDT (**Figure 5G**); this interesting observation was confirmed later in the UV-SCC model (see below).

**FIG. 5.**
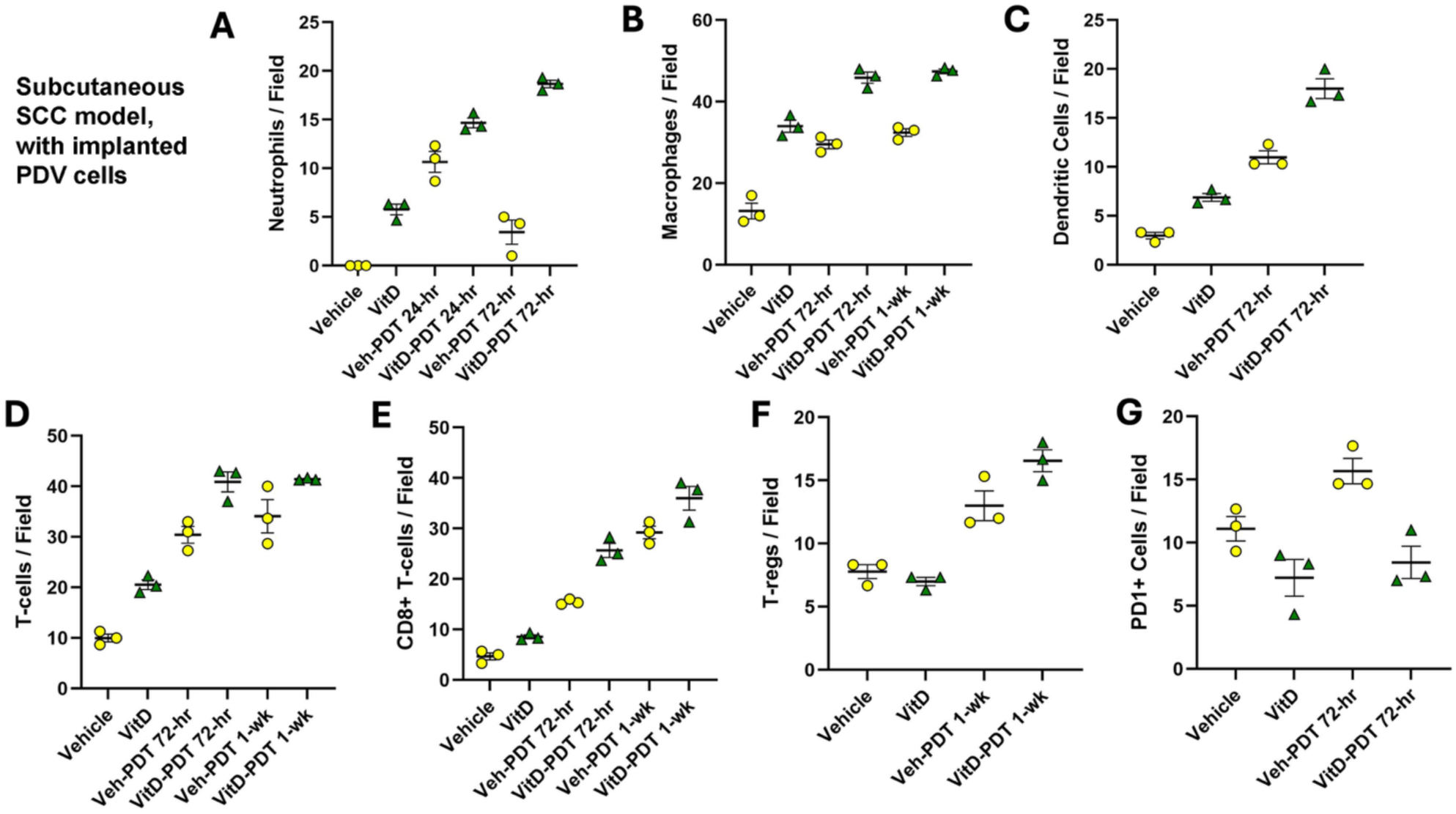
Recruitment of innate and adaptive immune cell into implanted PDVC57B SCC tumors following VitD ± PDT. Quantification of (**A**) neutrophils; (**B**) macrophages; (**C**) dendritic cells; (**D**) CD3+ (pan) T-cells; (**E**) CD8⁺ T-cells; (**F**) T-regs; and (**G**) PD1+ cells per field is shown. *n* = 3 mice per group. Each point represents the average of three fields per lesion from each mouse, with mean ± SEM displayed.

### Changes in innate immune cell populations in the peripheral blood after VitD±PDT

To compare local tumor immune cell recruitment with data on systemic changes, peripheral blood was collected at the same time as UV-SCC tumor harvest at 72 hr post-PDT (see **Figure 1**),and analyzed by flow cytometry. Results demonstrated that the proportion of dendritic cells (DCs) amongst total immunocytes (CD45^+^ cells) was significantly increased after PDT alone and after VitD+PDT, relative to vehicle (p = 0.03 for both); DCs also increased after VitD alone and after PDT or VitD+PDT at 1 week, but did not reach statistical significance (**Supplemental Figure 1A**). In contrast, the proportions of neutrophils (**Supplemental Figure 1B**) and monocytes (**Supplemental Figure 1C**) showed no significant differences across treatment groups.

### Changes in T-cell proportions and activation status in the peripheral blood after VitD ±PDT

The proportions of total T cells and of CD4⁺ or CD8^+^ T cells within the T-cell compartment were not significantly altered by treatment (**Supplemental Figure 2A-C**). However, T-regs were significantly reduced after PDT or VitD+PDT relative to vehicle alone (at 1 week, but not at 72 hr); no change was observed for VitD alone (**Figure 6B**). Activated CD69^+^ CD4⁺ T cells were significantly increased with VitD alone, PDT (72 hr, 1 week), or VitD+PDT (72 hr, 1 week) as compared with vehicle (**Figure 6C**).

**FIG. 6.**
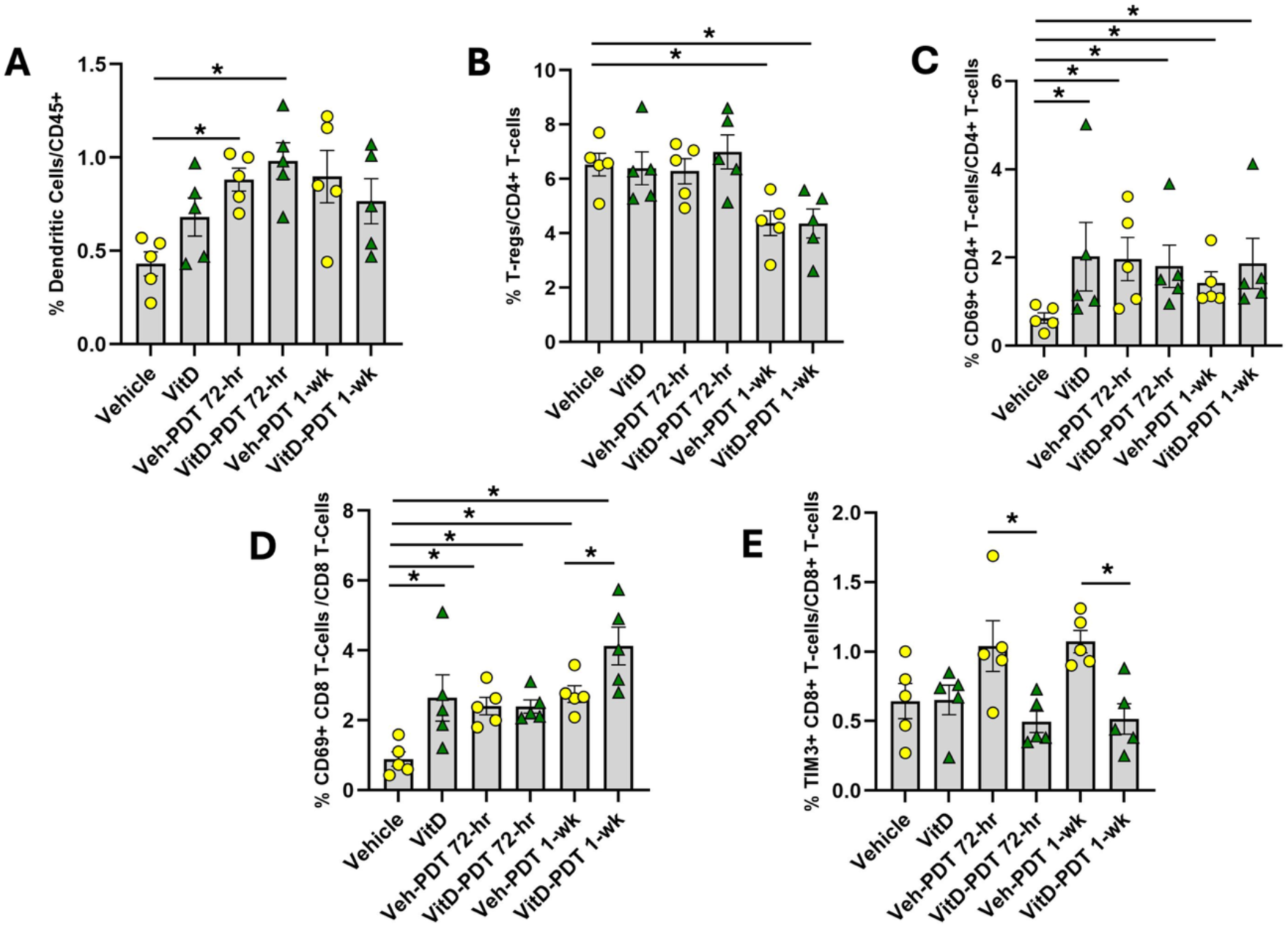
Changes in relative numbers of innate and adaptive immunocytes in the peripheral blood of UV-SCC mice. Cell numbers were determined by flow cytometry at 72 h after VitD ± PDT, relative to no-treatment controls. Only those cell types displaying significant changes are presented in this figure; see Online Supplement for data from all the cell types that were surveyed. (**A**) Dendritic cells, as proportion of all myeloid cells; (**B**) regulatory T-cells (Tregs; CD25⁺CD127⁻/CD4⁺ T-cells; (**C**) activated helper T = CD69+CD4+/CD4+ T cells; (**D**) activated cytotoxic T = CD69+CD8+/CD8+ T cells; (**E**) exhausted cytotoxic T = TIM3+CD8+/CD8+ T cells. *n* = 5 mice per group; points represent individual mice with mean ± SEM. Statistical analysis was performed as in Figure 2. (*), Significance level *p* < 0.05.

Activated CD69⁺ CD8⁺ T cells were increased significantly, relative to vehicle, after treatment with VitD alone, PDT (72 hr, 1 week), and VitD+PDT (72 hr, 1 week) (**Figure 6D**). At 1 week, the CD69⁺ CD8⁺ T cells were also significantly increased in VitD+PDT versus PDT alone (**Figure 6D**). TIM3⁺ CD8⁺ T-cells were modestly increased at 72 hr and 1-week post-PDT (not significant) compared with vehicle; however, the addition of VitD to PDT significantly reduced TIM3⁺ CD8⁺ T-cell proportions at both 72-hr and 1 week compared with PDT alone (**Figure 6E**). No significant differences were observed in naïve, central memory, effector memory T-cells (**Supplemental Figure 2D-F**) or in ICOS⁺ CD4⁺ or ICOS⁺ CD8⁺ T-cells (**Supplemental Figure 2G-H**) across treatment groups.

### Transcriptomic analysis reveals pro-tumorigenic signaling pathways affected differentially by Vitamin D and PDT

To investigate transcriptional changes induced by treatment, we performed bulk RNA sequencing on UV-SCC tumor tissue after exposure to vehicle, VitD alone, PDT alone, and VitD+PDT groups at 72 hr post-treatment. Gene set enrichment analysis (GSEA) results, in which the effects of each treatment relative to vehicle are expressed as a -fold change (Normalized Enrichment Score, NES), are listed in **Table 1** along with the statistical significance of each change.

**Table 1.** Gene Set Enrichment Analysis (GSEA) of Signaling Pathway Activation in Murine SCC Tumors treated with PDT alone, VD alone, or sequential VD+PDT.

| Pathway Gene Set<br>(Hallmark) |  | PDT<br>vs.<br>Vehicle | VD<br>vs.<br>Vehicle | VD + PDT<br>vs.<br>Vehicle | VD + PDT<br>vs.<br>PDT |
| --- | --- | --- | --- | --- | --- |
| Interferon alpha response | NES<br>*p* (*) | 1.93<br>7.2E-06 | 2.02<br>1.8E-07 | 3.02<br>2.5E-09 | 2.14<br>1.4E-08 |
| Interferon gamma response | NES<br>*p* | 1.63<br>1.6E-04 | 1.83<br>1.7E-07 | 2.75<br>2.5E-09 | 1.90<br>3.4E-04 |
| IL2 – Stat 5 signaling | NES<br>*p* | 1.30<br>0.033 | 1.62<br>8.6E-05 | 1.49<br>0.005 | 1.08<br>0.310 |
| TGF beta signaling | NES<br>*p* | 0.87<br>0.793 | 1.42<br>0.030 | -1.10<br>0.405 | -1.25<br>0.190 |
| Epithelial-to-mesenchymal transition (EMT) | NES<br>*p* | 1.50<br>0.003 | -1.90<br>2.6E-09 | -1.40<br>0.017 | -1.78<br>1.1E-06 |
| TNF alpha signaling via NFk-B | NES<br>*p* | 1.08<br>0.401 | 1.59<br>2.7E-04 | 1.01<br>0.560 | 1.04<br>0.429 |
| Angiogenesis | NES<br>*p* | -1.31<br>0.177 | -1.66<br>0.006 | -1.57<br>0.033 | -1.20<br>0.256 |

Murine UV-induced SCC tumors were harvested at 72 h after PDT; after VitD treatment alone (3 days of topical calcitriol); or a 72 h after a combination of VitD pretreatment + PDT. Values shown in the table: *NES*, normalized enrichment scores (-fold) of pairwise comparisons; *p*, adjusted P value (< 0.05 is significant). Increases (green) or decreases (red) in pathway activity are indicated. Yellow boxes highlight statistically significant changes.

GSEA revealed that PDT and VitD upregulated interferon signaling pathways compared with vehicle. Specifically, PDT significantly increased **interferon-α responses** vs. vehicle (NES = 1.93), as did VitD (NES = 2.02); the combination of VitD+PDT was even more effective at enhancing the interferon-α response (NES = 3.02). A similar pattern was seen for **interferon-γ responses**; PDT or VitD enhanced gene activity (NES = 1.63) while the VitD+PDT combination further amplified these effects (NES = 2.75). For both interferon pathways, activity scores for combination VitD+PDT were significantly higher than for PDT alone.

Several other pathways were also signficantly upregulated by PDT or VitD, but showed no interaction. **IL2–STAT5 signaling** was enriched by PDT and by VitD, yet the combination of VitD+PDT did not appear to raise the signal any higher than did the individual agents. **TGF-β signaling** was modestly enhanced by VitD, but showed no increase with PDT, and showed a trend toward downregulation when VitD and PDT were combined. **TNF-α signaling** was significantly enriched only by VitD, but not by VitD+PDT.

Conversely, two pathways were specifically downregulated in the presence of VitD. **Epithelial–mesenchymal transition (EMT)** was slightly increased after PDT (NES = 1.50), but decreased after VitD treatment (NES = -1.90), and was also decreased after combination VitD+PDT, suggesting that VitD has a dominant effect in suppressing EMT. **Angiogenesis** pathway genes were also signficantly suppressed by VitD, alone and together with PDT. PDT alone also suppressed angiogenesis, but the trend was not statistically significant.

### Vitamin D induces changes in the tumor microenvironment that favor anti-tumor immune stimulation

The large numerical increases in innate and adaptive immune cells recruited locally after VitD treatment seems likely to increase overall anti-tumoral immune functions, but further evidence is needed. In that regard, two changes in specific immune cell sutypes are indicators of likely improvement in functional outcome. First, the ratio of proinflammatory (M1) to quiescent (M2) macrophages was observed to change in the presence of VitD. Each type of macrophage (M1 CD86+ macrophages, **Figure 7A**; and M2 CD206+ macrophages, **Figure 7B**) was increased after VitD, PDT, or VitD+PDT, but to different degrees. As a result, the ratio of M1/M2 was reduced by PDT alone, yet maintained after VitD or VitD+PDT (**Figure 7C**). Thus, M1-mediated anti-tumor activity seems to be suppressed by PDT, but VitD prevents that undesirable side effect.

**FIG. 7.**
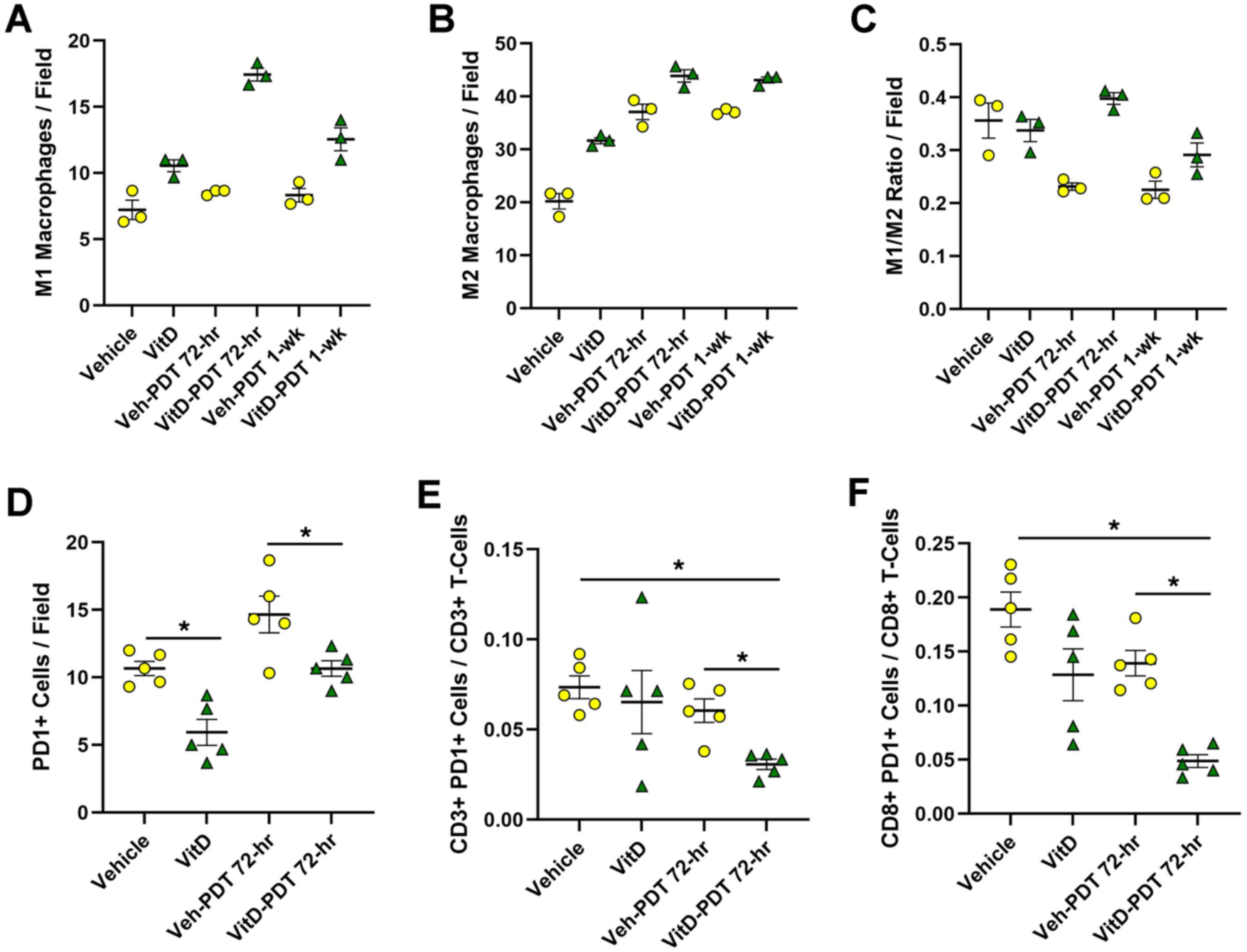
Vitamin D pretreatment induces changes in macrophages and in PD1-expressing T cells that are likely to improve overall anti-tumor immune action after PDT. Tissue sections from UV-SCC tumors were harvested at 72 hours after VitD ± PDT, and immunostained cells counted and quantified as shown in the graphs. *<u>Macrophage subsets</u>*: (**A**) M1 macrophages, CD86+; (**B**), M2 macrophages, CD206+; (**C**) M1/M2 macrophage ratio, *n* = 3 mice per group; each point represents the average of three fields per lesion from each mouse, with mean ± SEM displayed. *<u>PD1-expressing T-cell subsets</u>*: (**D**) PD1+ cells (**E**), CD3+PD1+ T-cells (**F**) and CD8+PD1+ T-cells. For D-F, there were *n* = 5 mice (lesions) per group, three fields per lesion averaged from each mouse, with mean ± SEM displayed. (*), statistical analysis using Kruskal–Wallis tests and Mann–Whitney U-tests for pairwise comparisons, *p* < 0.05.

Second, we observed that PD1+ (exhausted) T cell numbers were reduced in the presence of VitD relative to vehicle (**Figure 7D**). When examining specific T-cell subsets by double labeling, both the PD1+CD3+ T-cells (**Figure 7E**) and the cytotoxic PD1+CD8+ T-cells (**Figure 7F**) were significantly reduced by the VitD+PDT combination.

## DISCUSSION

In this study, we demonstrate that neoadjuvantal VitD in combination with ALA-based PDT reshapes the tumor immune landscape in murine SCC, producing additive and qualitatively distinct immune effects compared with either treatment alone.

Regarding immunogenic cell death, we observed robust upregulation of DAMPs (calreticulin and HMGB1) and simultaneously a local increase in neutrophils, macrophages, and dendritic cells. All of these effects occured after PDT or VitD alone, but were much stronger after combined VitD+PDT. Given the role of DAMPs in activating APCs[28] and priming adaptive immunity, these findings suggest that VitD amplifies the earliest immune-activating events initiated by PDT. Interestingly, PDT and VitD appeared to have selective and opposing effect upon macrophage subtypes. After PDT alone, M2 macrophages were selectively increased (2-fold), whereas M1 macrophages barely changed (∼1.2-fold) (**Fig. 7A-C**), representing a shift toward an immunosuppressive phenotype. However, adding VitD to the PDT regimen elevated M1 macrophagess at the expense of M2, boosting the M1/M2 ratio and creating a more favorable environment for anti-tumor activity (**Fig. 7A-C**). VitD appears to induce a qualitative reprogramming of macrophages that counters PDT-induced M2 expansion by greatly increasing M1 cells, thereby prolonging the anti-tumor immune response.

Within the adaptive immune compartment, robust recruitment of CD3+CD8+ (cytotoxic) T cells was observed, although functional benefits of this may be counteracted by increased FoxP3⁺ T-regs after PDT or VitD+PDT (**Fig. 4**). VitD reduced PD-1⁺ T-cells relative to vehicle controls (**Fig. 5G**), and VitD+PDT further reduced PD-1 expression relative to PDT alone (**Fig. 5G**, **Fig 7D-F**), suggesting that VitD may counteract checkpoint upregulation after PDT. The latter observation is consistent with prior research in non-small cell lung cancer (NSCLC)[29], where VitD was shown to decrease checkpoint marker expression on cytotoxic T cells.

Transcriptomic analysis (**Table 1**) provided functional context for changes in immune cell subsets observed histologically. GSEA revealed strong enrichment of interferon (IFN)-α responses (potentially linked to antigen presentation and innate immune activation[30]) and of IFN-γ responses (associated with cytotoxic T-cell effector function[30]) after PDT, VitD, and VitD+PDT. These changes align with findings of increased innate and adaptive immune cell infiltration, suggesting that recruited cells were not only more abundant but also functionally primed. TNF-α signaling (consistent with pro-inflammatory cytokine release and neutrophil recruitment[31]) was upregulated by VitD, in line with increases in intratumoral myeloid cells, while IL2–STAT5 signaling (critical for T-cell proliferation and survival[32]) was moderately enriched. VitD (alone or together with PDT) suppressed pro-tumorigenic EMT and angiogenesis pathways, suggesting additional pathways for limiting tumor progression.

Immune profiling of peripheral blood offered findings that supplement the tumor histological and transcriptional data. Circulating dendritic cells were increased by PDT and combination treatment, suggesting that enhanced systemic antigen presentation might occur in parallel with the increase in tumor-infiltrating dendritic cells. Peripheral T-regs were reduced by VitD+PDT despite being increased intratumorally, suggesting recruitment from circulation rather than systemic expansion. While this trafficking could represent a local counter-regulatory response to PDT-induced inflammation, the observed rise in circulating activated CD69⁺ T cells and decline in TIM3⁺ exhausted CD8⁺ T cells in the VitD+PDT group indicate that overall effector quality is improved, not blunted. While no major differences were observed across naïve, central memory, effector memory, and ICOS⁺ T-cell subsets, this may reflect the limited one-week observation period. Fnally, although PD-1 was included in the flow cytometry panel, minimal PD-1 expression was observed in peripheral blood. This finding is consistent with prior literature demonstrating that PD-1 expression is predominantly localized to tumor-infiltrating lymphocytes rather than peripheral blood T cells[33].

In the broader context of VitD effects upon immunity (which are typically contradictory and context dependent), VitD can be immunosuppressive in chronic inflammatory conditions where it downregulates pro-inflammatory cytokines and enhances IL-10 and T-reg function, thereby reducing tissue damage[34–36], reinforcing tolerance, and inhibiting development of autoimmune diseases[34, 37, 38]. However, VitD can also be immunostimulatory, as during infections where VitD enhances innate antimicrobial responses via cathelicidins, defensins, and PRR activation[39]. In cancer, VitD engages VDR-dependent transcription to inhibit tumor proliferation, angiogenesis, and tumor survival[40, 41]. VitD shifts the local microenvironment toward an effector-dominant state[40, 42], increasing NK and CD8⁺ T cell activity in glioblastoma and breast cancer [43, 44], enhancing Th1 cytokine production and reducing immune checkpoint activity in NSCLC[29], increasing CD8⁺ T cell density in colorectal cancer[45], and improving survival via strengthened immune signatures in melanoma[46]. In the therapeutic context of PDT, VitD acts through multiple mechanisms including immune activation (these data), enhanced PpIX photosensitizer accumulation[47], and altered cytokine secretion by cancer-associated fibroblasts[48].

This study has some limitations. First, neutrophils and monocytes in our FACS studies were defined only by surface markers, preventing their distinction from myeloid derived suppressor cells (PMN-MDSCs or M-MDSCs). However, the transcriptomic data indicate a predominantly pro-inflammatory state with reduced angiogenesis; since MDSCs typically promote immunosuppression and angiogenesis, the immune cells observed here are unlikely to represent suppressor subsets. Second, our UV-SCC model (chosen to mimic field cancerization and human SCC development) can be quite variable in terms of tumor size and morphology. To partially address this, a more-uniform SCC model (PDV) was studied in parallel and found to engender similar immune effects as the UV-SCC tumors. Third, our RNA-seq analyses were limited to 72 hr post-PDT, a timepoint that captures innate and early adaptive responses but may miss later changes. Finally, we were unable to quantify CD4⁺ T cells by immunofluorescence due to technical challenges. In future studies we hope to address the above limitations and conduct functional experiments to ask whether VitD+PDT can synergize with immune checkpoint blockade to improve tumor resolution. Future experiments will also address whether anti-EMT and anti-angiogenesis activities (suggested by our transcriptomic data) can lower rates of metastases after VitD+PDT in a murine tumor model. Ultimately, the VitD+PDT combination should be investigated in human clinical trials of SCC for its potential to improve therapuetic outcomes and serve as an immune-priming strategy.

In summary, data in this report collectively show that VitD enhances PDT immune effects by amplifying the magnitude of immune activation and qualitatively reprogramming it. The combination treatment promotes IFN-driven cytotoxic responses, increases M1 macrophages, reduces inhibitory checkpoints (PD-1, TIM3), suppresses EMT and angiogenesis, and shifts the immune landscape toward more effective cell-mediated immunity.

## Data Availability Statement

All data supporting the findings of this study are available within the paper and its Supplementary Information.

## Author Contribution Information

Alan Shen, Sanjay Anand, Marcela Diaz, Tayyaba Hasan, and Edward Maytin contributed to the study conception and design. Material preparation, data collection and analysis were performed by Sanjay Anand, Cheng-En Cheng, Alan Shen, Benjamin Kovacic, and Jennifer Powers. The first draft of the manuscript was written by Alan S. Shen and all authors commented on previous versions of the manuscript. All authors read and approved the final manuscript.”

