## Supplemental Figs 1 and 2 for "Photodynamic priming with Vitamin D and ALA-based PDT induces intratumoral immune cell recruitment and signaling pathway activation in murine cutaneous squamous cell carcinoma"

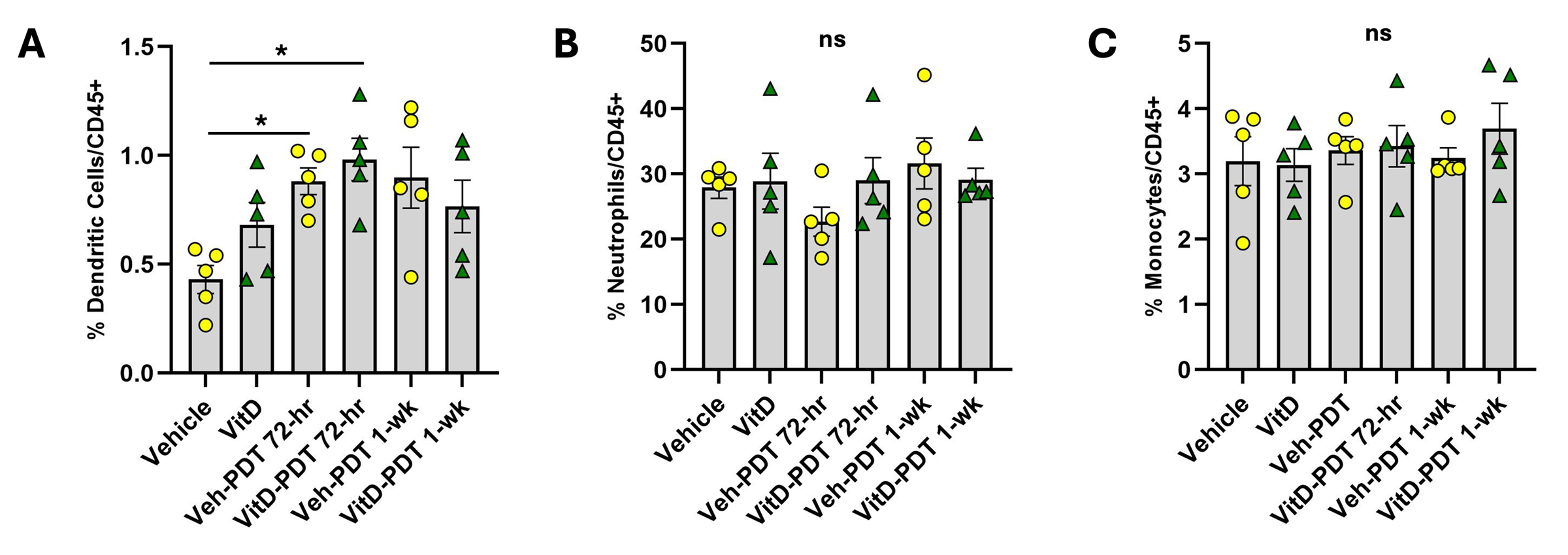

**Supplemental Fig. 1. Flow cytometry analysis of innate immune cells in peripheral blood from UV-induced SCC mice following VitD ± PDT.** Flow cytometric quantification of dendritic cells (**A**), neutrophils (**B**), and monocytes (**C**) as a percentage of total immune cells (CD45⁺) at 72 hours and 1-week post-treatment. *n* = 5 mice per group; points represent individual mice with mean ± SEM. Statistical analysis used Kruskal–Wallis for overall group comparisons, and when significant, Mann–Whitney U-tests for pairwise comparisons with adjustment for multiple comparisons utilizing the Benjamini-Hochberg correction. Significance levels: (*****), *p* < 0.05; ***ns,*** not significant (*p* ≥ 0.05).

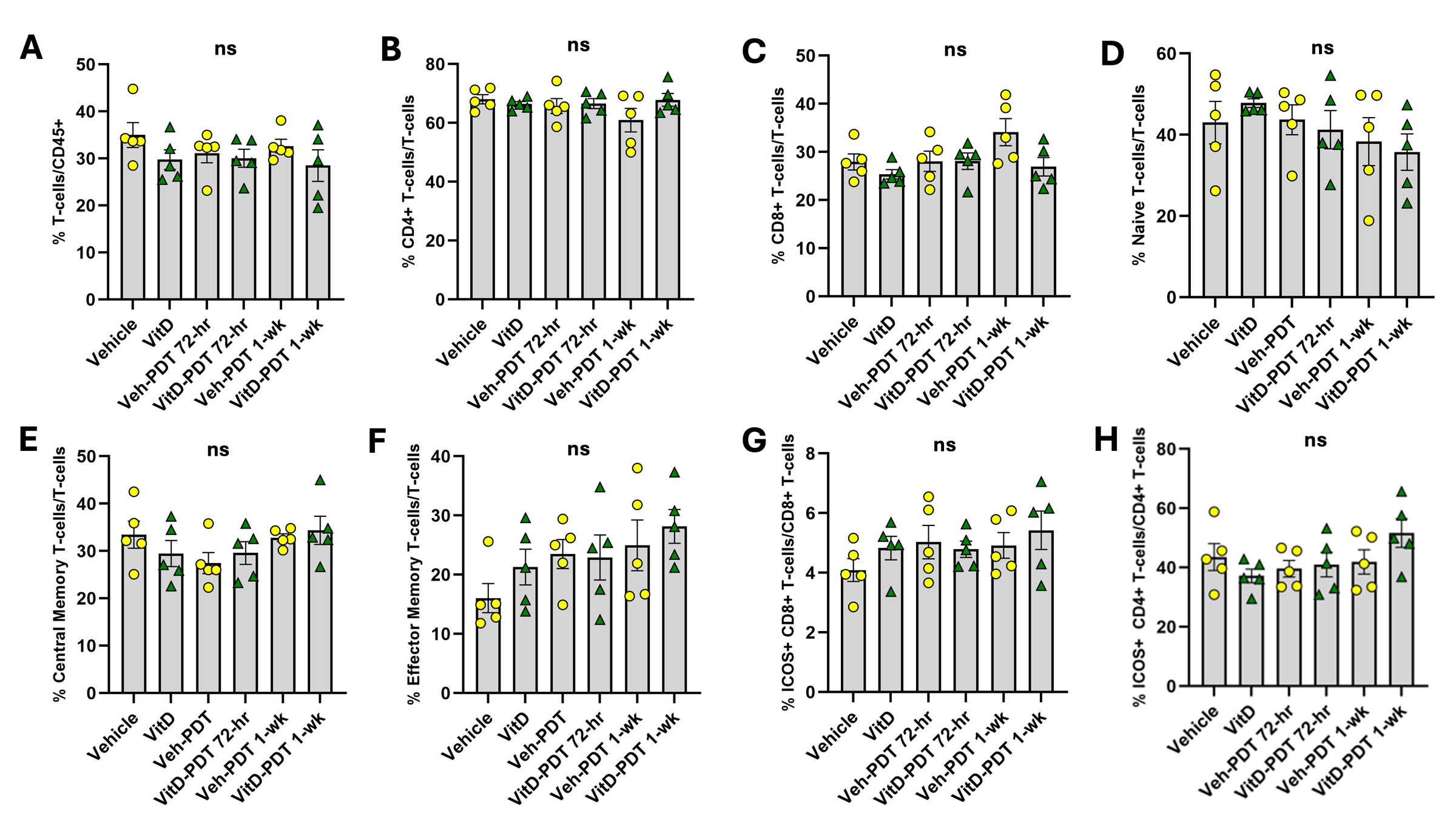

**Supplemental Fig. 2.** **Flow cytometry analysis of T-cell subsets in peripheral blood from UV-induced SCC mice following VitD ± PDT.** Flow cytometric quantification of (A) total T-cells out of CD45⁺ immune cells, (B) CD4⁺ T-cells out of total T-cells, (C) CD8⁺ T-cells out of total T-cells, (D) naïve T-cells out of total T-cells, (E) central memory T-cells out of total T-cells, (F) effector memory T-cells out of total T-cells, (G) ICOS+ CD8+ T-cells out of CD8+ T-cells, and (H) ICOS+ CD4+ T-cells out of CD4+ T-cells. *n* = 5 mice per group; points represent individual mice with mean ± SEM. Statistical analysis performed as in Suppl. Fig. 1. Significance levels: (*****), *p* < 0.05; ***ns,*** not significant (*p* ≥ 0.05).
